# The effect of autopolyploidy on population genetic signals of hard sweeps

**DOI:** 10.1101/753491

**Authors:** Patrick Monnahan, Yaniv Brandvain

## Abstract

Searching for population genomic signals left behind by positive selection is a major focus of evolutionary biology, particularly as sequencing technologies develop and costs decline. The effect of the number of chromosome copies (i.e. ploidy) on the manifestation of these signals remains an outstanding question, despite a wide appreciation of ploidy being a fundamental parameter governing numerous biological processes. We clarify the principal forces governing the differential manifestation and persistence of the signal of selection by separating the effects of polyploidy on rates of fixation versus rates of diversity (i.e. mutation and recombination) with a set of coalescent simulations. We explore what the major consequences of polyploidy, such as a more localized signal, greater dependence on dominance, and longer persistence of the signal following fixation, mean for within- and across-ploidy inference on the strength and prevalence of selective sweeps. As genomic advances continue to open doors for interrogating natural systems, studies such as this aid our ability to anticipate, interpret, and compare data across ploidy levels.

## Intro

Whole genome duplication (i.e. polyploidization) is common, particularly within the plant kingdom (1). This macromutation can impact both macroevolutionary processes of colonization, speciation, and extinction (2), and the microevolutionary processes of mutation, drift and natural selection (3). Modern sequencing technologies provide insight into how selection shapes the genomic landscape of divergence and could allow researchers to test hypotheses about how polyploidy alters the tempo and mode of adaptation. However, before we can analyze sequence data to test such hypotheses, we must understand how the same selective pressures alter our ability to recognize a sweep in polyploids.

Numerous features of polyploids could impact the tempo and mode of adaptation. For example, all else equal, an increase in chromosome number will increase the mutational target size, and thus, the rate of adaptation in a mutation-limited world (4). The additional set of chromosomes may also change dominance and epistatic relationships, potentially offering novel routes to adaptation (5). Finally, the transition to polyploidy can shock the genomic system, initiating manifold selective pressures (6). With the ongoing development of sequencing and computational tools, researchers will surely turn to population genomic studies to better understand the evolutionary consequences of polyploidy and to address hypotheses concerning the nature of adaptation in polyploids.

However, most of our scans for selection have been developed with diploids in mind, which hinders population genetic analysis of selection in natural polyploids. Thus, while we would like to know if polyploidy fundamentally changes the action of selection, we first need to know how two inherent features of polyploidy -- larger population mutation and recombination rates, due to larger effective chromosome number, and slower responses to selection due to lower variance in fitness -- change the neutral variation around adaptive substitutions, and subsequently change our power to identify and localize sweeps. Importantly, the effect of ploidy on each of these factors is to promote the retention of neutral diversity in polyploids, although they likely differ in their qualitative and quantitative effects. In contrast to the more straightforward, ‘factor-of-two’ effects of polyploidy on mutation and recombination, the effect on selection is more complicated due to the added potential for an allele to be masked or diluted in heterozygous genotypes, effectively dampening the single-locus selection response across much of the range of dominance conditions and allele frequency (9).

With appropriate consideration, coalescent simulations can generate haplotype data for an arbitrary ploidy level (7), and thus guide interpretation of patterns of sequence variation. The implicit assumption of free recombination among haplotypes, restricts our analysis to autopolyploids (where chromosomal copies derive from a single ancestral species), which, by some estimates are as or more frequent than allopolyploids (resulting from hybridization) (8). We utilize these simulations to disentangle how the multifarious effects of chromosome number impact the population genomic signals left behind by selection, again, focusing on the relative impact on mutation and recombination versus the effects on selection and fixation times.

We show that the effects of selection on neutral diversity are both more striking and more locally-restricted in polyploids as compared to diploids. Further analyses reveal that the limited reach of selective sweeps in polyploids is primarily attributable to the extended fixation time, rather than their increased population recombination rates. The differential effects of ploidy on the selection signal are also highly dependent on the dominance of the mutation, particularly for recessive cases. Lastly, we find that the signal of selection persists for more or less time in higher ploidies, depending on the particular metric used. In sum, we highlight the many ways that chromosome number fundamentally alters the selective process and the consequences that this has for population genomic inference and comparison across ploidy levels.

## Methods

### Simulating polymorphism

We simulate selective sweeps with *mssel* (10), under the standard (*ceteris paribus*) assumption that polyploidy simply increases the number of chromosome copies, *c*_*s*_ (7). Thus, the coalescent time unit equals *c*_*s*_ *\* N*, and the population mutation and recombination rates equal *θ = 2Nc*_*s*_*μL* and ρ*=2Nc*_*s*_*rL*, respectively, where *L* is the length of sequence to simulate (1 Mb for all simulations; see Table 1 for a list of parameter descriptions). By modelling ploidy with the scalar, *c*_*s*_, we assume no preferential pairing between homologs and no “double reduction.” We further assume an equal number of individuals, *n*, sampled for all ploidies (we set *n* = 10 for all analyses, thus the number of chromosomes sampled scales with ploidy).

**Table 1.** Summary of parameters used and referred to throughout the text.

| Abbv | Parameter | Description | Values |
| --- | --- | --- | --- |
| $L$ | Sequence Length | Number of base pairs of sequence to simulate | 1 Mb |
| $H$ | Dominance Scalar | determines relationship between individual allele frequency and fitness | (0.1, 0.3, 0.5, 0.6, 0.9) |
| $g_1$ | Fuse Generation | Generations prior to start of sweep that the two populations fused | (1, $10^3$, $10^4$) |
| $g_2$ | Sample Generation | Generations since fixation of beneficial allele | (1, $10^3$, $10^4$) |
| $\mu$ | Mutation Rate | Mutation rate; per-base, per-generation | $10^{-8}$ |
| $N$ | Population Size | Number of individuals in a population | ($10^3$, $10^4$) |
| $r$ | Recombination Rate | probability of recombination event; per-base, per-generation | $10^{-8}$ |
| $s$ | Selection Coefficient | difference in fitness between alternative homozygotes | (0.01, 0.1) |
| $c_s$ | Simulation Ploidy | ploidy for determining population parameters ($\theta$, $\rho$, etc.) | (2, 4, 8) |
| $c_t$ | Trajectory Ploidy | ploidy for generating sweep trajectory | (2, 4, 8) |

We simulate a simple demographic scenario in which an ancestral population splits in two at time *g*_*1*_, at which point a beneficial mutation arises in the middle of the chromosome (at basepair 500,000) in one population and ultimately fixes. We sample *n * c*_s_ haplotypes from each population (Supp. Figure 1) *g*_*2*_ generations after a sweep is completed. We use the non-selected population as a neutral baseline and for the calculation of between-population measures of selection (F_ST_, XP-EHH, etc.).

### Generating allele frequency trajectories

We developed a ploidy-general form of the allele frequency recursion equation as:

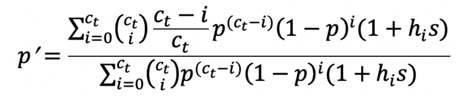

where *c*_*t*_ is ploidy number (diploids=2, tetraploids=4, etc.), *i* is the number of non-mutant alleles, *h*_*i*_ is the dominance coefficient for the genotype with *i* out of *c*_*t*_ non-mutant alleles, *p* is the allele frequency, and *s* is the selection coefficient. When *c*_*t*_ = 2, we recover the familiar formula for allele frequency change in diploids.

To compare across ploidies, we assume that the selection coefficient for a beneficial mutation is the difference in relative fitness between alternative homozygotes, and that the dominance coefficient for a heterozygous genotype is a simple function of the frequency of beneficial alleles within the individual, and 3) this function is constant across ploidy levels. We use a single value, which we call the *dominance scalar* (*H*), to control the form of the dominance function (Figure 1C; similar to (4)). For a particular *H* (−0.5 < *H* < 0.5), a genotype’s dominance coefficient is calculated as 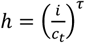, where τ = ∣ (10 *H*)^*sgn*(*H*)^ ∣. With additivity (*H* = 0), the relationship is linear, whereas a negative (recessive) value of *H* results in τ > 1, and thus a convex function, and a positive (dominant) value results in a concave function (i.e. 0 < τ < 1).

**Figure 1.**
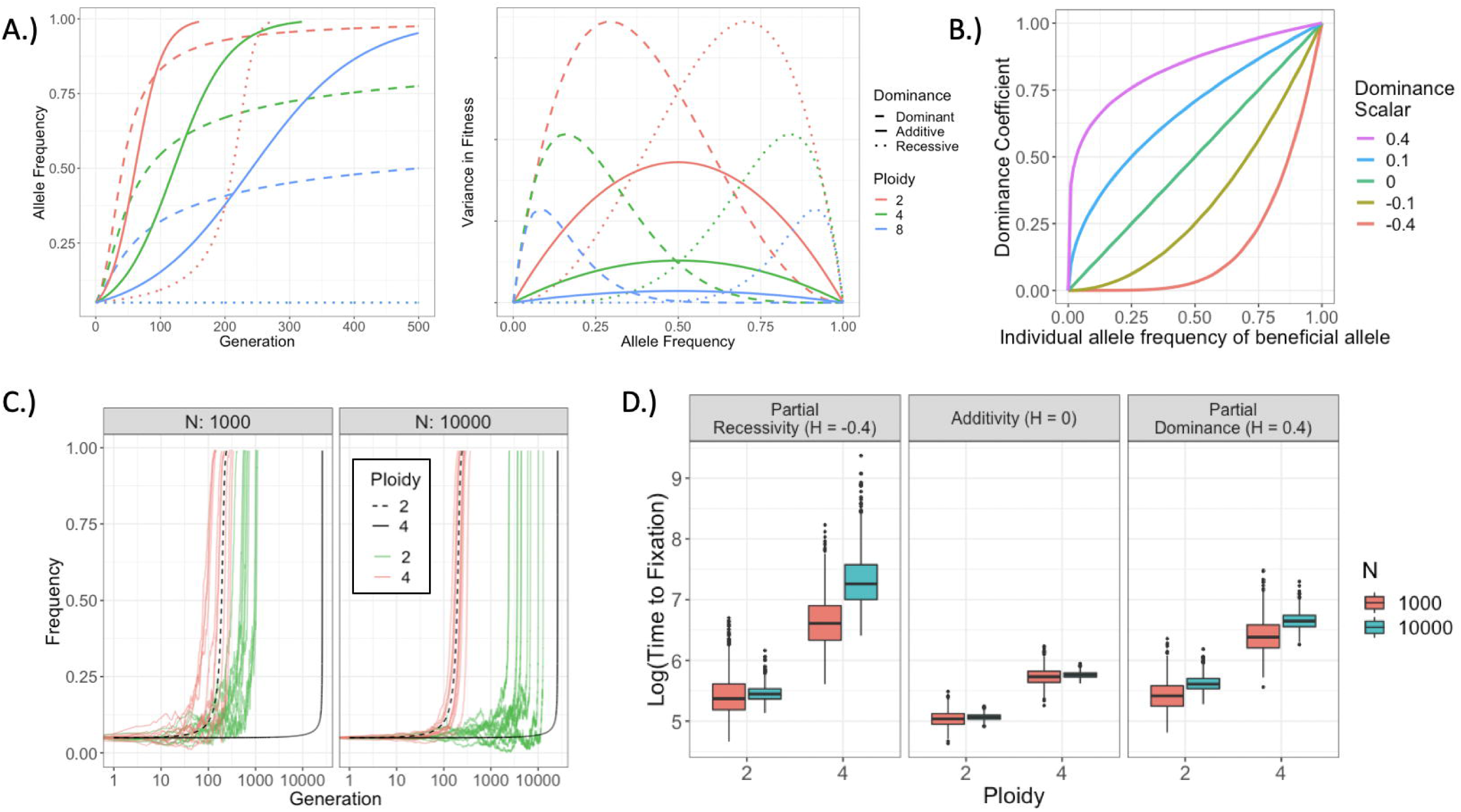
A) Left, Slower rates of allele frequency change in higher ploidies. Right, reduced variance in fitness in higher ploidies at intermediate allele frequencies. s=0.1. B) Dominance scheme to determine dominance coefficients based on allele frequency within the given genotype. C) Allele frequency trajectories for recessive mutations at different population sizes (N). The black lines illustrate the expectation with infinite population size. D) Effects of population size on fixation time for different dominance scenarios.

We generated stochastic allele frequency trajectories as a series of sampling events from the binomial distribution:

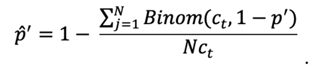

iterating until the beneficial was fixed or lost. Because we are only interested in successful sweeps, we conditioned on fixation by only recording trajectories in which the beneficial allele was fixed.

### Analysis

The numerous metrics designed to detect selection vary widely in their assumptions and robustness to various confounding forces, such as demography. To make valid comparisons across ploidy, we are restricted to metrics which do not implicitly or explicitly assume diploidy. We chose two common metrics based on nucleotide diversity (Tajima’s D and F_ST_) and two haplotype-based metrics (iHS and XPEHH). We calculated pairwise nucleotide diversity (*π*), F_ST_, and Tajima’s D in overlapping windows (step size of ½ full window size) using the R package *PopGenome* (12). After experimenting, we found that 1 Mb / (*N /* 50) windows captured meaningful variation in these summaries across all parameters that we investigated. We calculated iHS (13) and XP-EHH (14) with the R package, *rehh* (15), using the *parse_ms()* function from msr (https://github.com/vsbuffalo/msr) to file format conversion.

## Results & Discussion

### Effect of ploidy on patterns of diversity following a selective sweep

The effect of selection on neutral diversity is stronger, but more locally restricted in higher ploidies, whereas it is locally weaker, but more diffuse in diploids (Figure 2A). Assuming that the rate of adaptation does not differ by ploidy, a greater proportion of the genome will be affected by linked selection in lower ploidies. In higher ploidies, the greater magnitude of the reduction in diversity was largely the result of the fact that higher ploidies have a higher baseline level of diversity, a result that is made clear by the strong effect of *c*_*s*_ on the magnitude of a sweep (Figure 2B). The reduced breadth of the dips in higher ploidies was largely driven by the elevated time to fixation, and less so due to the increased recombination rate, as can be seen by the dominant effect of *c*_*t*_ (“trajectory ploidy”) on sweep breadth (although recombination rate also clearly has a marginal effect on breadth; Figure 2B). Together, this suggests that, with whole genome sequencing, sweeps will be better localized with higher ploidies, but with reduced representation sequencing, sweeps are more likely to be missed with higher ploides.

**Figure 2.**
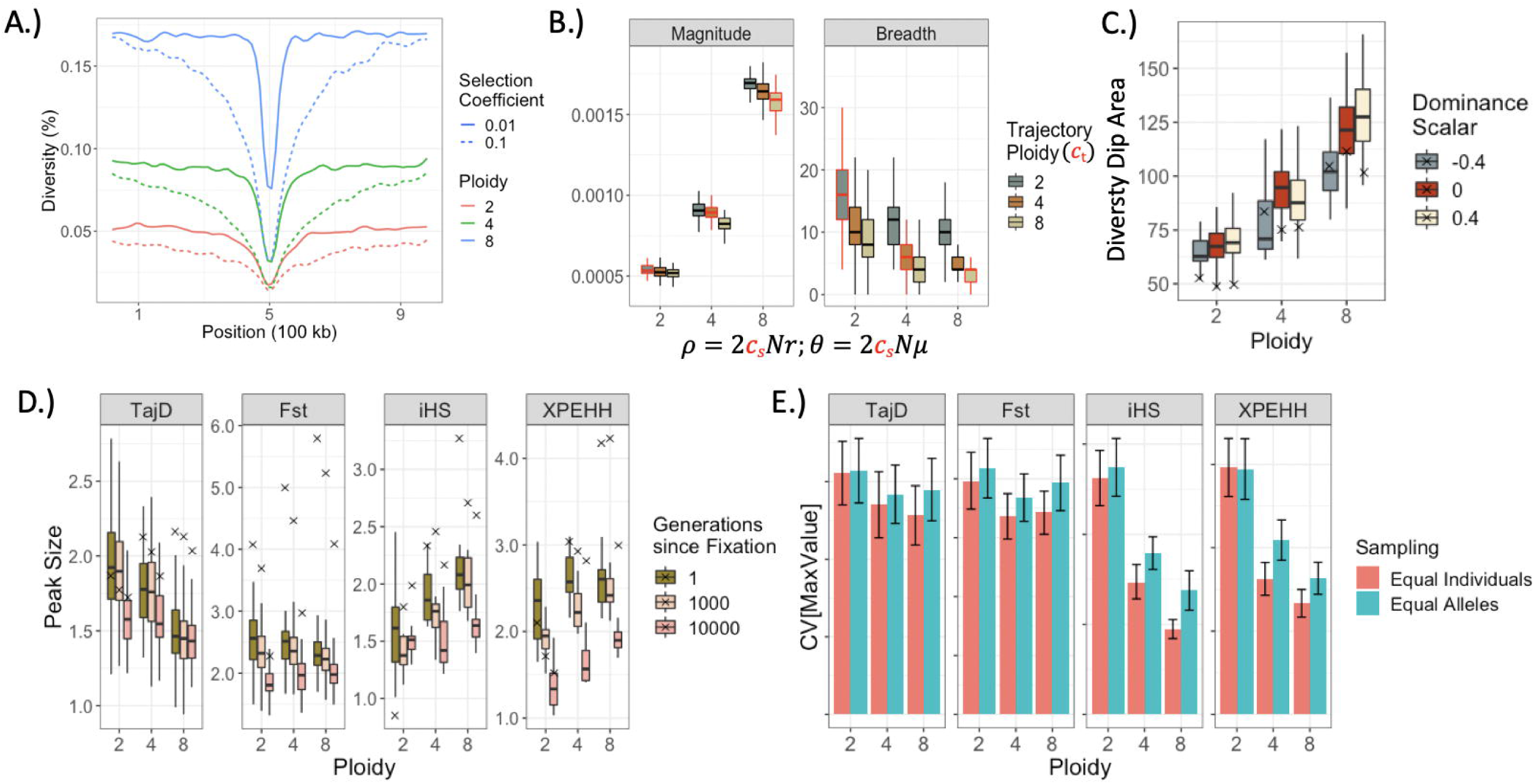
A.) Smoothed profile of diversity (20 replicates) observed at different ploidy levels for strong and weak selection. B.) Relative impacts of diversity/recombination and fixation rates on profiles of diversity. C.) Ploidy-dependent effects of dominance on diversity. X denotes median for N=1,000. D.) Effect of ploidy on the persistence of the selection signal. H=0. Fst is standardized within replicates. X denotes median maximum value across replicates, and bars are the integrated area under the curve. E.) Coefficient of variation for the median (across replicates) of the maximum value observed for each replicate. Bars are estimates from 1000 replicates, and error bars are from resampling 200 replicates 1000 times. For all plots, N=10,000; s=0.01.

We also find that ploidy interacts with dominance to modulate the manifestation of sweep signals. In contrast to an additive locus, where there is an approximate factor-of-two effect of ploidy on fixation time, differences can be much greater for non-additive mutations, particularly when the mutation is recessive (Figure 1A, C-D). For example, with strong recessivity (*H* = −0.4) fixation time increases by ∼10x whenever ploidy is doubled. This results in a reduced signal of non-additive mutations in higher ploidies, whereas the effect of dominance is minimal in diploids. In other words, recessive mutations are not only more likely to be lost in higher ploidies, but they may also be more likely to go undetected when not lost (Figure 2C; Supp. Figure 2).

This ploidy-dependent effect of dominance is partly mitigated in smaller populations. In large populations, allele frequency change due to selection can be much slower at certain allele frequencies in higher ploidies, as a consequence of the fact that genotypes with the greatest differences in relative fitness are very rare (16) (Figure 1A, C-D). For recessive mutations, the slowdown occurs initially because the mutant homozygote bearing the full manifestation of selection is much rarer in higher ploidies (*p*^*2*^, *p*^*4*^, and *p*^*8*^ for dip-, tetra-, and octaploids, respectively). Although greater stochasticity in allele frequency change in small populations will result in more frequent loss of these alleles, for those that survive, it may push the frequency above the critical threshold in which selection can gain traction (Figure 1C). Once past this point, selection acts very quickly on recessive mutations, occasionally resulting in greater effects on diversity than dominant or additive mutations, in contrast to what is observed in larger populations (see “X” points on Figure 2C for median values when *N* = 1000). Dominant mutations, on the other hand, stall at more intermediate frequencies, following sharp increases in frequency early on. Here, drift can interfere with the weakened efficacy of selection and ultimately produce a weaker signal in genomic data.

### Measures of selection

Ploidy modestly amplified evidence for a sweep for most metrics of selection, particularly for the maximum observed values for the haplotype-based statistics (i.e. XPEHH and iHS; Figure 2D, see ‘X’ points for max values). The exception was for total peak area of Tajima’s D, which showed a weaker signal with higher ploidy, and standardized Fst, which showed no effect. The variance of these selection metrics also tends to decrease in higher ploidies (Figure 2E), which is, again, particularly pronounced for the haplotype-based statistics. The reduced variance is not simply a consequence of sampling a greater number of chromosomes from the population (for a given number of individuals). Even when sampling an equivalent number of alleles from each population, there is a notable reduction in variance in higher ploidies (Figure 2E). Thus for equivalent sampling/sequencing effort, many metrics of selection are more precisely estimated with higher plody.

We also find that with elevated ploidy, signals of selection persist for a longer time, as it takes longer to reach mutation drift equilibrium with increasing coalescent times. This was the case for the majority of the metrics that we investigated, with the exception of the iHS statistic. The irregular iHS pattern in diploids results from a preponderance of undefined values when taking ratios of haplotype scores in populations fixed for a single haplotype. Interestingly, despite the increased effective recombination rate in polyploids, haplotype-based measures of selection persist longer, again reflecting their slower return to equilibrium. Recent development in phasing algorithms, necessary for calculating haplotype scores, will thus greatly advance our ability to detect recent selection in autopolyploids (17).

## Conclusion

Polyploidy fundamentally changes microevolutionary processes. While the increased mutational opportunity in polyploids may boost adaptation over evolutionary timescales (4), the increased sojourn time of the beneficial allele has important consequences for shaping genomic diversity, and thus evolutionary inference. Understanding these changes and their consequences is important, as natural polyploids are increasingly being interrogated with modern sequencing methods. In demonstrating the inherent effects of ploidy on particular population genomic measures, our (and future) work will guide in the identification and interpretation of signals of selection with higher ploidy.

## Acknowledgments

We thank Arthur Zwanepoel for help deriving the recursion equation.

