## Supplemental Figures for "The effect of autopolyploidy on population genetic signals of hard sweeps"

Supplemental Figure 1. Coalescent simulation framework

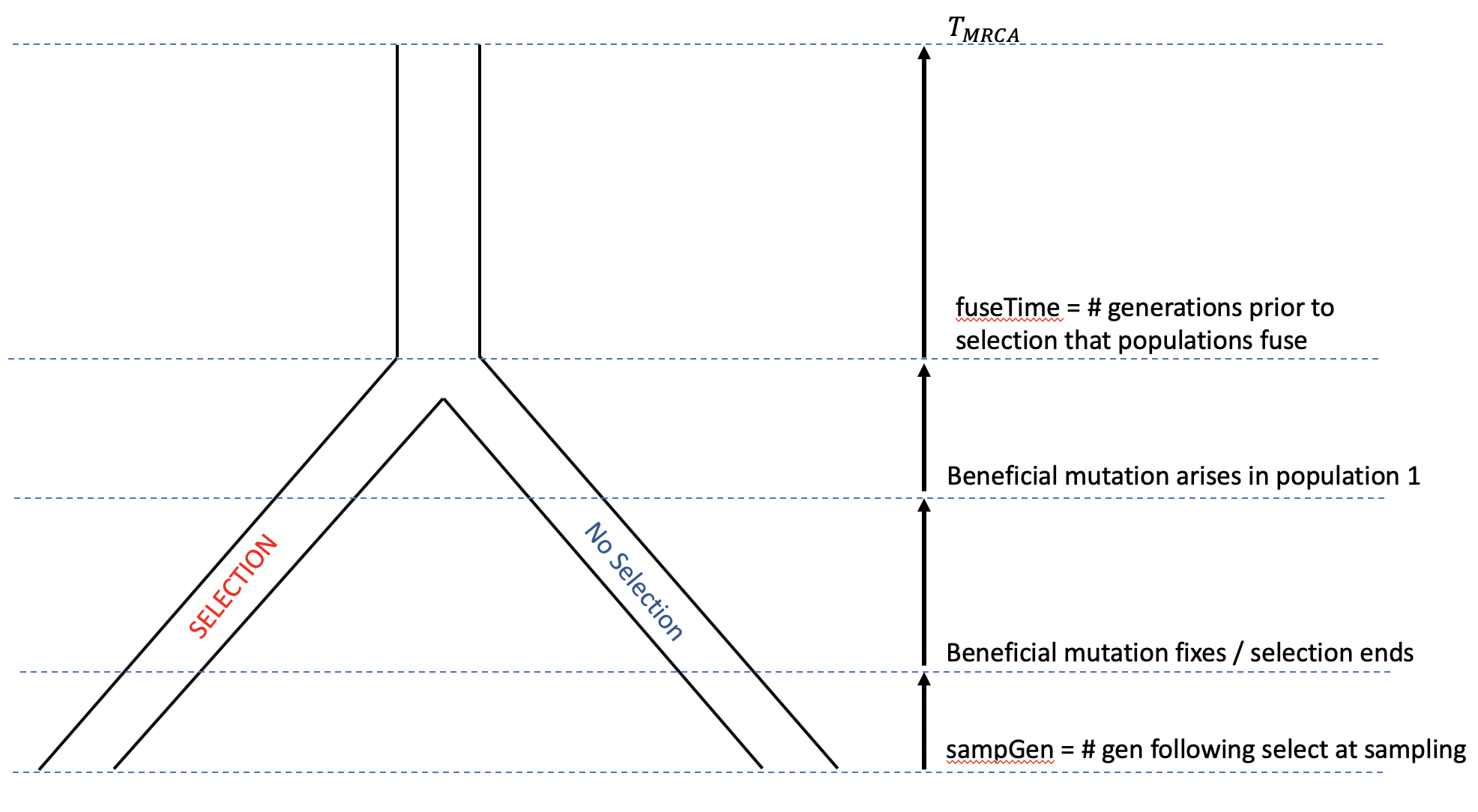

Supplemental Figure 2. Effect of dominance on various metrics of selection

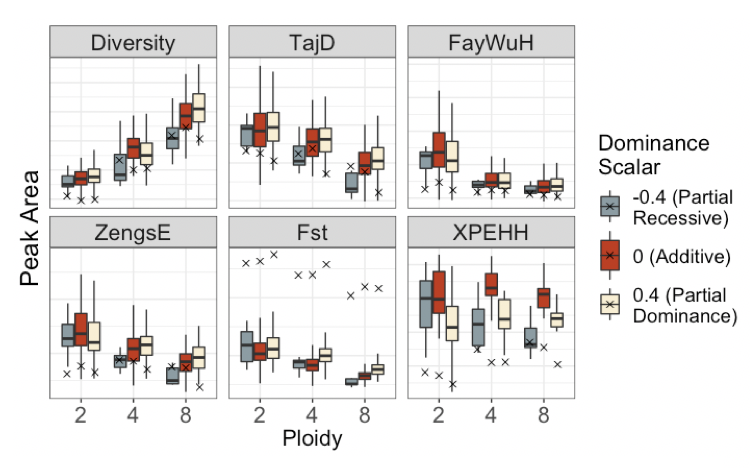
